## Supplementary material for "Analyzing heterogeneity in Alzheimer Disease using multimodal normative modeling on imaging-based ATN biomarkers": Table 1

### **Table 1**: Descriptive statistics for the ADNI-ADS and ADRC-ADS datasets. Statistical differences were assessed using two-sided ANOVA (continuous variables) and chi-squared tests (categorical. variables). Significant p-values are highlighted in bold with *: 0.01 < p < 0.05, **: 0.005 < p < 0.01, ***: p < 0.001. Abbreviations: SD = standard deviation, ANOVA = analysis of variance, CDR = Clinical Dementia Rating, MMSE = Mini-Mental State Examination.

|  | ADNI-ADS | ADRC-ADS | p-value |
| --- | --- | --- | --- |
| N | 231 | 129 | - |
| Sex, Male: Female | 108:123 | 48:81 | **p = 0.035*** |
| Age (mean +/- SD) | 73.6 +/- 6.9 | 71.5 +/- 8.3 | **p = 0.006**** |
| CDR (0/0.5/>=1) | 121/80/30 | 98/24/7 | **p < 0.001***** |
| MMSE (mean +/- SD) | 24.5 +/- 3.2 | 26.5 +/ 3.7 | **p < 0.001 ***** |
