## Supplementary material for "Analyzing heterogeneity in Alzheimer Disease using multimodal normative modeling on imaging-based ATN biomarkers": Table 2

### **Table 2**: Comparison between DSI across all modalities (DSI_all), and the ATN summary metrics (hippocampal volumes, amyloid burden, and tau index) with respect to association with the composite cognitive scores (memory, executive functioning, and language). β represents the slope and p represents the p-value for linear regression, adjusted for age and sex. r represents the Pearson correlation coefficient.

|  | **Cognitive domain** | **ADNI-ADS** | | | **ADRC-ADS** | | |
| --- | --- | --- | --- | --- | --- | --- | --- |
|  |  | **β** | **p** | **r** | **β** | **p** | **r** |
| **DSI_all** | **Memory** | - 0.65 | p < 0.001 | **- 0.62** | - 0.71 | p < 0.001 | **- 0.68** |
|  | **Executive** | - 0.46 | p < 0.001 | **- 0.54** | - 0.52 | p < 0.001 | **- 0.56** |
|  | **Language** | - 0.39 | p < 0.001 | **- 0.47** | - 0.36 | p < 0.001 | **- 0.41** |
