## Supplementary methods and results for "Analyzing heterogeneity in Alzheimer Disease using multimodal normative modeling on imaging-based ATN biomarkers"

### Supplementary Methods (SM)

#### **SM1: MRI imaging in ADRC**

##### SM1.1 Image acquisition

MRI data were acquired on a Siemens Biograph mMR or Trio 3T scanner. Anatomical scans were acquired with a T1-weighted magnetization-prepared rapidly acquired gradient echo (MPRAGE) sequence. Images were acquired in a sagittal orientation with the following parameters: repetition time of 2,300 milliseconds, an echo time of 2.95 milliseconds, a flip angle of 9°, 176 slices, an in-plane resolution of 240 × 256, and a slice thickness of 1.2 mm.

##### SM1.2 Image preprocessing

Images were processed using FreeSurfer[1] (version 5.3) (<http://surfer.nmr.mgh.harvard.edu/>) for volumetric segmentation into Regions of Interest (ROI). The FreeSurfer processing pipeline included the following steps: (a) motion correction and segmentation of the subcortical white matter and deep gray matter volumetric structures on a T1 weighted image, (b) intensity normalization (c) registration to a spherical atlas which utilizes individual cortical folding patterns to match cortical geometry across participants and finally (d) parcellation of the cerebral cortex based upon the Desikan atlas. For all images failing a quality control check, the processing pipeline was rerun to ensure consistency across all scans.

#### **SM2: PET imaging in ADRC**

##### SM2.1 Image acquisition

Amyloid PET imaging was performed with [18F]-Florbetapir (FBP) and was acquired on a Biograph mMR (Siemens Medical Solutions, Malvern, PA). Participants who underwent FBP imaging received a 370 MBq dose of 18F-florbetapir, followed by a 20-minute scan captured 50 minutes post-injection. Tau PET imaging used the tracer [18F]-Flortaucipir (FTP) and was acquired on a Biograph 40 PET/CT scanner (Siemens Medical Solutions). Participants who underwent FTP imaging were administered an intravenous injection of [18F]-Flortaucipir (370 MBq dose), followed by a 20-minute scan captured 80 minutes post-injection.

##### SM2.2 Image preprocessing

All PET data from Knight ADRC were processed with an in-house pipeline using regions of interest derived from FreeSurfer (PET Unified Pipeline, <https://github.com/ysu001/PUP>). Initially, images were smoothed to achieve a target resolution of 8 mm FWHM (full width and half maximum) and underwent motion correction. Subsequently, the images were co-registered with the participant's T1weighted-MRI acquisition from the corresponding visit. The co-registered T1-weighted images were processed using FreeSurfer (version 5.3) for volumetric segmentation into ROIs. Amyloid FBP deposition was quantified with the average across the left and right lateral orbitofrontal, medial orbitofrontal, rostral middle frontal, superior frontal, superior temporal, middle temporal, and precuneus regions to derive average standardized uptake value ratios (SUVRs) in ROIs.[2] Likewise, tau FTP deposition was summarized with the average of bilateral entorhinal cortex, amygdala, inferior temporal lobe, and lateral occipital cortex to derive average SUVRs in ROIs.[3]

#### **SM3: Neuropsychological composites**

Following previous work[4], we computed neuropsychological composites for memory, executive functioning, and language. However, some modifications were added to the composites to account for availability of cognition data in ADNI and Knight ADRC. The specific tests from both datasets were calculated as follows: (i) **Memory**: The memory composite for ADNI consisted of the immediate and delayed recall items from both the Logical Memory IIa test[5] and the Rey Auditory Verbal Learning Test.[6] For Knight ADRC, the memory composite differed from ADNI and consisted of the delayed item recall from the Logical Memory IIa test[5], the Free and Cued Selective Reminding Test[7], and Associate Memory[5] (ii) **Executive functioning**: the executive function composite consisted of Trails Making Test Parts A & B[8] for both ADNI and Knight ADRC. (iii) **Language**: the language composite for both datasets included the category fluency for animals and vegetables[9], the Boston Naming Test [10], and the Multilingual Naming Test.[11] We converted all individual test scores into Z-scores relative to the respective control group (i.e., ADNI-CU and ADRC-CU) with low scores indicating more impairment. Next, we averaged the rescaled scores to compute the neuropsychological composite scores.

#### **SM4: Multimodal Normative Modelling**

##### SM4.1 Joint distribution between multiple modalities (mmVAE)

Our proposed mmVAE has separate modality-specific encoders and decoders for individual modalities. The main idea is to assume that the joint distribution over the multiple modalities factorizes into a product of single- modality data-generating distributions when conditioned on the latent space.[12,13] This assumption is used to derive the structure and factorization of the variational posterior. Without loss of generality, we assume that we have M modalities x_1_,x_2_ . . . , x_M_ ,, which are conditionally independent given the common latent variable z.[3] So, we assume a generative model of the form $p_{\theta}\left( x_{1},..x_{M},z \right)=p\left( z \right)\prod_{i=1}^{M} p_{\theta}\left( x_{i} | z \right)$. The conditional independence assumptions in the generative model imply a relation among the joint-modality posterior $p\left( z | x_{1},..,x_{M} \right)$ and the single-modality posteriors $p\left( z | x_{i} \right)$ as shown below.

$$p\left( z | x_{1},..,x_{M} \right)=\frac{p\left( x_{1},..,x_{M} | z \right)*p\left( z \right)}{p\left( x_{1},..,x_{M} \right)}=\frac{p\left( z \right)}{p\left( x_{1},..,x_{M} \right)}*\prod_{i=1}^{M} p\left( x_{i} | z \right)$$

$$p\left( z | x_{1},..,x_{M} \right)=\frac{p\left( z \right)}{p\left( x_{1},..x_{M} \right)}*\prod_{i=1}^{M} \frac{p\left( z | x_{i} \right)p\left( x_{i} \right)}{p\left( z \right)}$$

$$p\left( z | x_{1},..,x_{M} \right)=\frac{\prod_{i=1}^{M} p\left( z | x_{i} \right)}{\prod_{i=1}^{M-1} p\left( z \right)}*\frac{\prod_{i=1}^{M} p\left( x_{i} \right)}{p\left( x_{1},..x_{M} \right)}$$

$$p\left( z | x_{1},..,x_{M} \right)\propto\frac{\prod_{i=1}^{M} p\left( z | x_{i} \right)}{\prod_{i=1}^{M-1} p\left( z \right)}$$

We see that the joint-modality posterior $p\left( z | x_{1},..,x_{M} \right)$ is a product of individual single-modality posteriors $p\left( z | x_{i} \right)$, with an additional quotient by the prior $p\left( z \right)$. If we approximate $p\left( z | x_{i} \right)$ ≡ $q\left( z | x_{i} \right)p\left( z \right)$ where $q\left( z | x_{i} \right)$ the underlying inference network for each modality, we can avoid the quotient term $p\left( z \right)$. Now, we can approximate the joint posterior as shown below (Equation 1):

$$p\left( z | x_{1},..,x_{M} \right)\propto\frac{\prod_{i=1}^{M} p\left( z | x_{i} \right)}{\prod_{i=1}^{M-1} p\left( z \right)}\equiv\frac{\prod_{i=1}^{M} \left[ q\left( z | x_{1} \right)p\left( z \right) \right]}{\prod_{i=1}^{M-1} p\left( z \right)}=p\left( z \right)\prod_{i=1}^{M-1} q\left( z | x_{i} \right)$$

The joint modality posterior $p\left( z | x_{1},..,x_{M} \right)$represents the joint distribution between the multiple modalities. This is called the Product-of-Experts (POE) approach where we approximate the distribution of the joint posterior $p\left( z | x_{1},..,x_{M} \right)$ by the product of a prior-expert $p\left( z \right)$ and the approximate modality-specific posterior $q^{\left( z | x_{i} \right)}.$ This product distribution shown in Equation 1 is not solvable in closed form. However, if we approximate both $p\left( z \right)$and $q^{'\left( z | x_{i} \right)}$as Gaussian, then a product of Gaussian experts is itself Gaussian with mean μ and variance σ (Equation 3) where μ_i_ and σ_i_ are parameters of the i-th Gaussian expert and $T_{i}$ = σ^−1^.[12,14]

##### SM4.2 Model architecture

Our proposed multimodal normative modeling framework (mmVAE) had specific encoders and decoders for individual modalities. Cortical and subcortical brain volumes extracted from T1-weighted MRI, amyloid FBP and tau FTP scans (FreeSurfer) were used as input to three modality specific encoders. The latent space parameters of the individual modalities were combined by the Product of Experts (POE) layer to form the shared latent space (SM 3.1). The joint latent space was which passed through the modality-specific decoders for reconstructions.

##### SM4.3 Training details

**Conditioning on covariates**: We conditioned our proposed mmVAE on the age and sex of patients, to ensure that the deviations in regional brain volumes reflect only disease pathology and not deviations due to effects of covariates. Both age and sex were transformed into one-hot encoding vectors. After this transformation, each subject had an age vector with 44 positions, where each position corresponds to a year within the range of 47–91 years. In this vector, all positions have value zero except the one that indicates the subject’s age which has a value equal to 1. The subject’s sex was represented in a one-hot encoded vector with two positions, one for male and one for female. Both the modality-specific decoders used these vectors together with the latent code to reconstruct the brain data.

**Model hyperparameters**: mmVAE was trained using the Adam optimizer with model hyperparameters as follows: epochs = 500, learning rate = 10^−5^, batch size = 256 and latent dimension = 64. The encoder and decoder networks consisted of four fully connected layers of sizes 512, 256, 128, 64 and 64, 128, 256, 512, respectively.

### Supplementary References

[1]	Fischl B. FreeSurfer. Neuroimage 2012;62:774–81.

[2]	Su Y, D’Angelo GM, Vlassenko AG, Zhou G, Snyder AZ, Marcus DS, et al. Quantitative analysis of PiB-PET with freesurfer ROIs. PloS One 2013;8:e73377.

[3]	Mishra S, Gordon BA, Su Y, Christensen J, Friedrichsen K, Jackson K, et al. AV-1451 PET imaging of tau pathology in preclinical Alzheimer disease: defining a summary measure. Neuroimage 2017;161:171–8.

[4]	Earnest T, Bani A, Ha SM, Hobbs DA, Kothapalli D, Yang B, et al. Data‐driven decomposition and staging of flortaucipir uptake in Alzheimer’s disease. Alzheimer’s & Dementia 2024.

[5]	Wechsler D. Wechsler memory scale-revised. Psychological Corporation 1987.

[6]	Rey A. L’examen clinique en psychologie. 1958.

[7]	Grober E, Buschke H. Genuine memory deficits in dementia. Developmental Neuropsychology 1987;3:13–36.

[8]	Reitan RM. Validity of the Trail Making Test as an indicator of organic brain damage. Perceptual and Motor Skills 1958;8:271–6.

[9]	Gladsjo JA, Schuman CC, Evans JD, Peavy GM, Miller SW, Heaton RK. Norms for letter and category fluency: demographic corrections for age, education, and ethnicity. Assessment 1999;6:147–78.

[10]	Kaplan E, Goodglass H, Weintraub S. Boston naming test 2001.

[11]	Gollan TH, Weissberger GH, Runnqvist E, Montoya RI, Cera CM. Self-ratings of spoken language dominance: A Multilingual Naming Test (MINT) and preliminary norms for young and aging Spanish–English bilinguals. Bilingualism: Language and Cognition 2012;15:594–615.

[12]	Kumar S, Payne PR, Sotiras A. Normative modeling using multimodal variational autoencoders to identify abnormal brain volume deviations in Alzheimer’s disease. Medical Imaging 2023: Computer-Aided Diagnosis, vol. 12465, SPIE; 2023, p. 1246503.

[13]	Wu M, Goodman N. Multimodal generative models for scalable weakly-supervised learning. Advances in Neural Information Processing Systems 2018;31.

[14]	Cao Y, Fleet DJ. Generalized product of experts for automatic and principled fusion of Gaussian process predictions. arXiv Preprint arXiv:14107827 2014.

### Supplementary Results

#### **Supplementary Tables**

#### **Table S1**: Descriptive statistics for the ADNI-CU and ADRC-CU datasets. Statistical differences were assessed using two-sided ANOVA (continuous variables) and chi-squared tests (categorical. variables) respectively. Significant p-values are highlighted in bold with *: 0.01 < p < 0.05, **: 0.005 < p < 0.01, ***: p < 0.001. Daggers within cells indicate significant differences between the CU & ADS cohorts for the same source dataset († p < 0.001) (see Table 1). Abbreviations: SD = standard deviation, ANOVA = analysis of variance, CDR = Clinical Dementia Rating, MMSE = Mini-Mental State Examination.

|  | ADNI-CU | ADRC-CU | p-value |
| --- | --- | --- | --- |
| N (%) | 434 | 301 | - |
| Sex, Male: Female | 205:229 **(†)** | 127:174 **(†)** | **p = 0.026*** |
| Age (mean +/- SD) | 71.5 +/- 7.2 **(†)** | 68.2 +/- 7.3 **(†)** | **p < 0.001***** |
| CDR (0/0.5/>=1) | 434/0/0 | 301/0/0 | - |
| MMSE (mean +/- SD) | 29.1 +/ 1.3 **(†)** | 29.5 +/ 1.7 **(†)** | p = 0.067 |

**Table S2:** Number of regions with abnormal (statistically significant) deviations across MRI, amyloid and tau for both ADNI-ADS and Knight ADRC-ADS.

| Modality | Group comparison | Number of abnormal regions (**ADNI**) | Number of abnormal regions (**ADRC**) |
| --- | --- | --- | --- |
| MRI | mild or severe dementia vs CU  very mild dementia vs CU  preclinical AD vs CU | 56  22  0 | 50  25  0 |
| Amyloid | mild or severe dementia vs CU  very mild dementia vs CU  preclinical AD vs CU | 84  75  85 | 81  77  82 |
| Tau | mild or severe dementia vs CU  very mild dementia vs CU  preclinical AD vs CU | 80  62  0 | 74  52  0 |

#### **Table S3**: Association between DSI calculated for individual modalities (DSI_mri, DSI_amyloid, DSI_tau) and the neuropsychological composite scores (memory, executive functioning, and language) for both ADNI-ADS and ADRC-ADS. β and p represent the slope and p-value for linear regression, adjusted for age and sex. r represents the Pearson correlation coefficient.

| DSI | Composite scores | ADNI-ADS | | | ADRC-ADS | | |
| --- | --- | --- | --- | --- | --- | --- | --- |
|  |  | β | p | r | β | p | r |
| DSI_mri | Memory | - 0.05 | p < 0.001 | - 0.34 | - 0.12 | p < 0.001 | - 0.42 |
|  | Executive | - 0.02 | p < 0.001 | - 0.25 | - 0.025 | p < 0.001 | - 0.24 |
|  | Language | - 0.03 | p < 0.001 | - 0.27 | - 0.045 | p < 0.001 | - 0.29 |
| DSI_amyloid | Memory | - 0.54 | p < 0.001 | - 0.32 | - 0.48 | p < 0.001 | - 0.62 |
|  | Executive | - 0.36 | p < 0.001 | - 0.25 | - 0.44 | p < 0.001 | - 0.41 |
|  | Language | - 0.33 | p < 0.001 | - 0.21 | - 0.39 | p < 0.001 | - 0.39 |
| DSI_tau | Memory | - 1.1 | p < 0.001 | - 0.51 | - 0.85 | p < 0.001 | - 0.49 |
|  | Executive | - 0.79 | p < 0.001 | - 0.46 | - 0.85 | p < 0.001 | - 0.51 |
|  | Language | - 0.73 | p < 0.001 | - 0.37 | - 0.72 | p < 0.001 | - 0.39 |

#### **Table S4**: Post-hoc comparisons for CDR survival analysis in ADNI-ADS and ADRC-ADS (see Figure 6). Pairwise differences were calculated using long-ranked tests. P-values were False Discovery Rate (FDR) corrected. Significant p-values are marked in bold with *: 0.01 < p < 0.05, **: 0.005 < p < 0.01, ***: p < 0.001.

| comparison | p-value (ADNI-ADS) | p-value (ADRC-ADS) |
| --- | --- | --- |
| q1 vs q2 | **p = 0.018*** | p = 0.072 |
| q1 vs q3 | **p = 0.084** | **p = 0.0085**** |
| q1 vs q4 | **p < 0.001***** | **p < 0.001***** |
| q2 vs q3 | **p = 0.0065 **** | **p = 0.022 *** |
| q2 vs q4 | **p < 0.001***** | **p < 0.001***** |
| q3 vs q4 | **p < 0.001***** | **p < 0.001***** |

#### **Supplementary Figures**

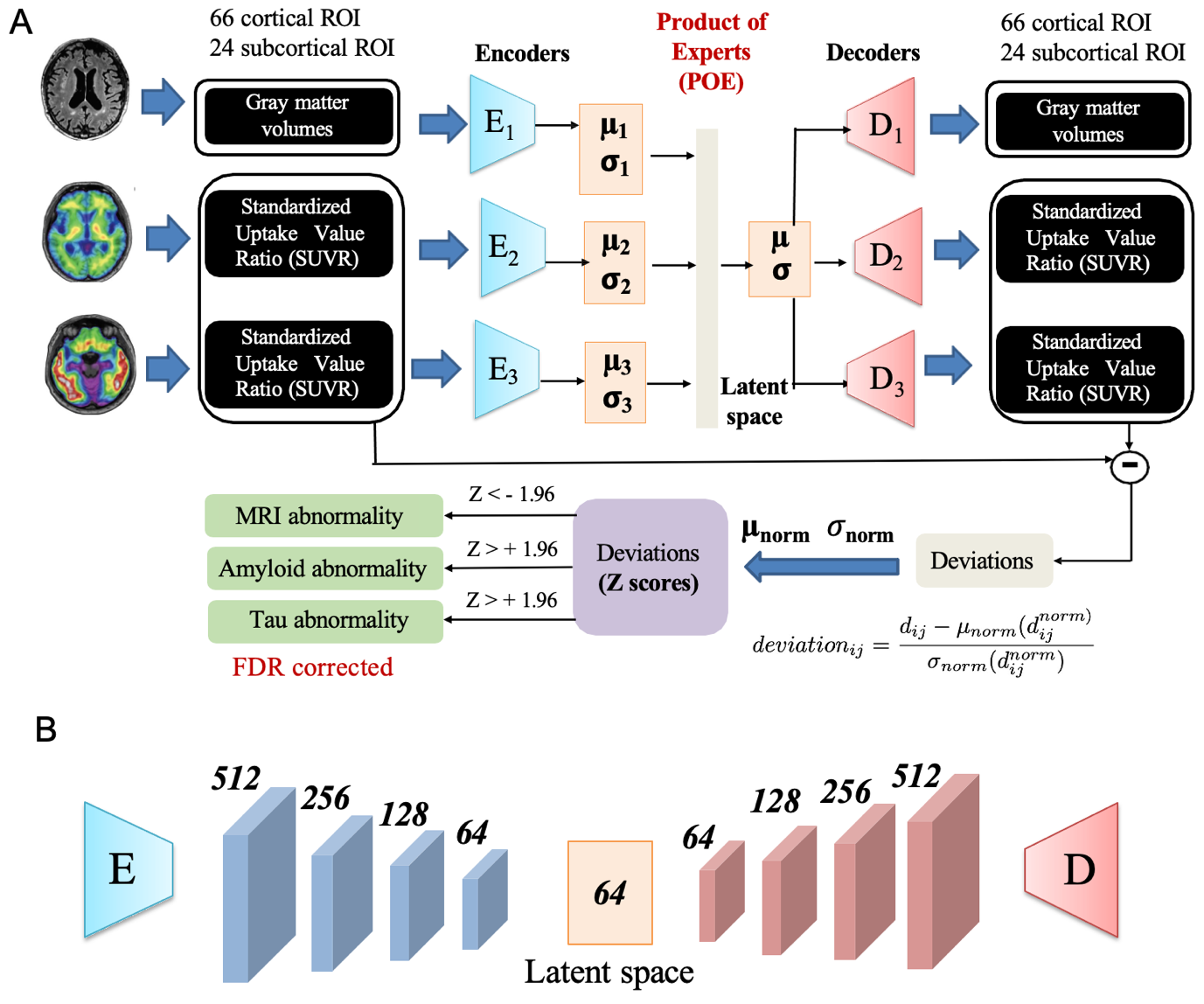

#### **Figure S1**: (**A**) Our proposed multimodal normative modeling framework (mmVAE). Cortical and subcortical brain volumes extracted from T1-weighted MRI, amyloid FBP and tau FTP scans (FreeSurfer) were used as input to three modality specific encoders. The latent space parameters of the individual modalities were combined by the Product of Experts (POE) layer to form the shared latent space. The joint latent space was which passed through the modality-specific decoders for reconstructions. The deviations for each modality were calculated by the reconstruction error between input and reconstructed data, normalized with respect to the CU (cognitively unimpaired) group to form Z-scores. The regions with abnormal (statistically significant) deviations were identified for each modality after FDR correction (Z < -1.96 for MRI and Z > 1.96 for amyloid and tau). (**B**) Structure of the encoder and decoder used in our analyses. The encoder and decoder networks have 4 fully connected layers of sizes 512, 256, 128, 64 and 64, 128, 256, 512, respectively, with a latent dimension of 64.

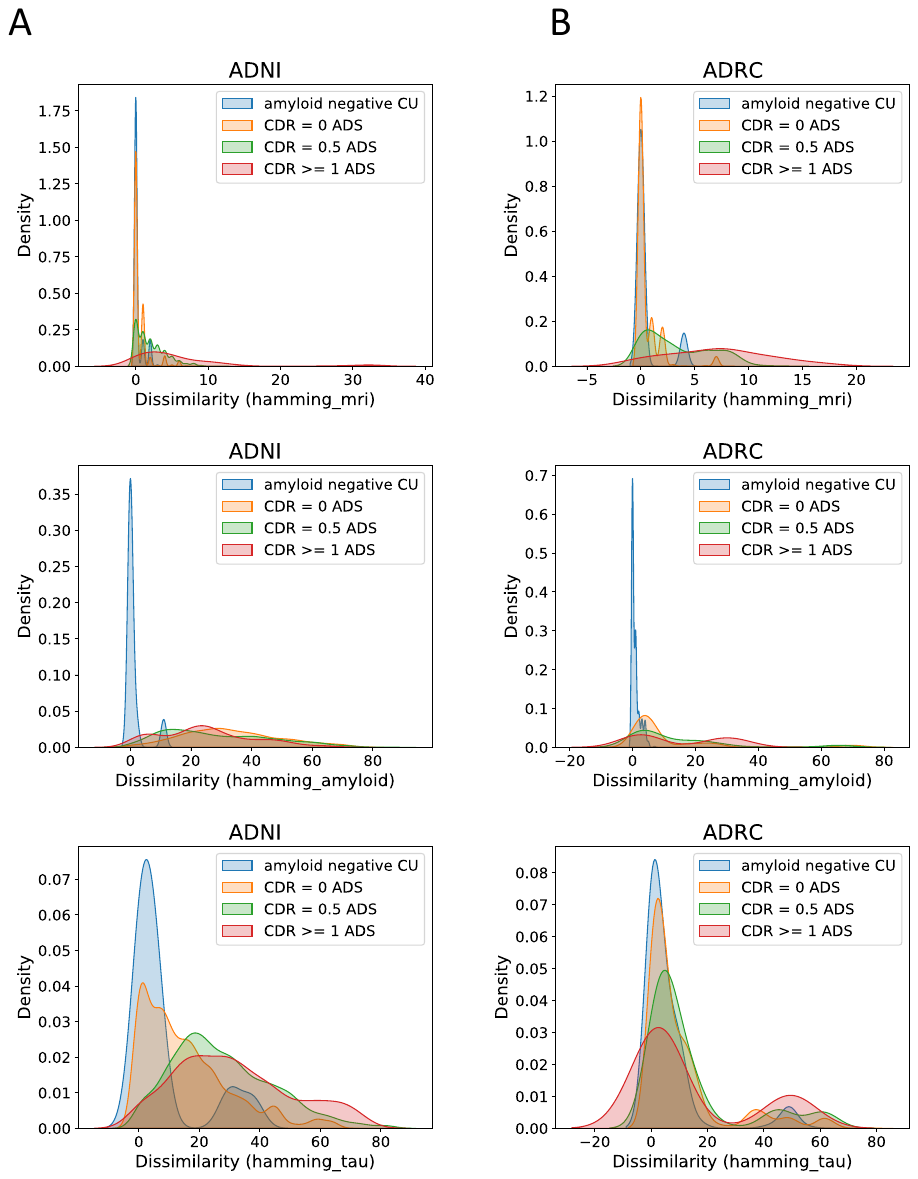

#### **Figure S2**: Hamming distance density (KDE plot) which illustrates the spread of dissimilarity in abnormality (calculated by Hamming distance) within each CDR group for ADNI (**S2A**) and Knight ADRC (**S2B**). The figures from top to bottom represent hamming distances calculated from abnormal deviations across mri, amyloid and tau respectively (hamming_mri, hamming_amyloid, hamming_tau see Section 2.5.4). Higher hamming distance values indicated more heterogeneity in abnormality patters even within a particular group.

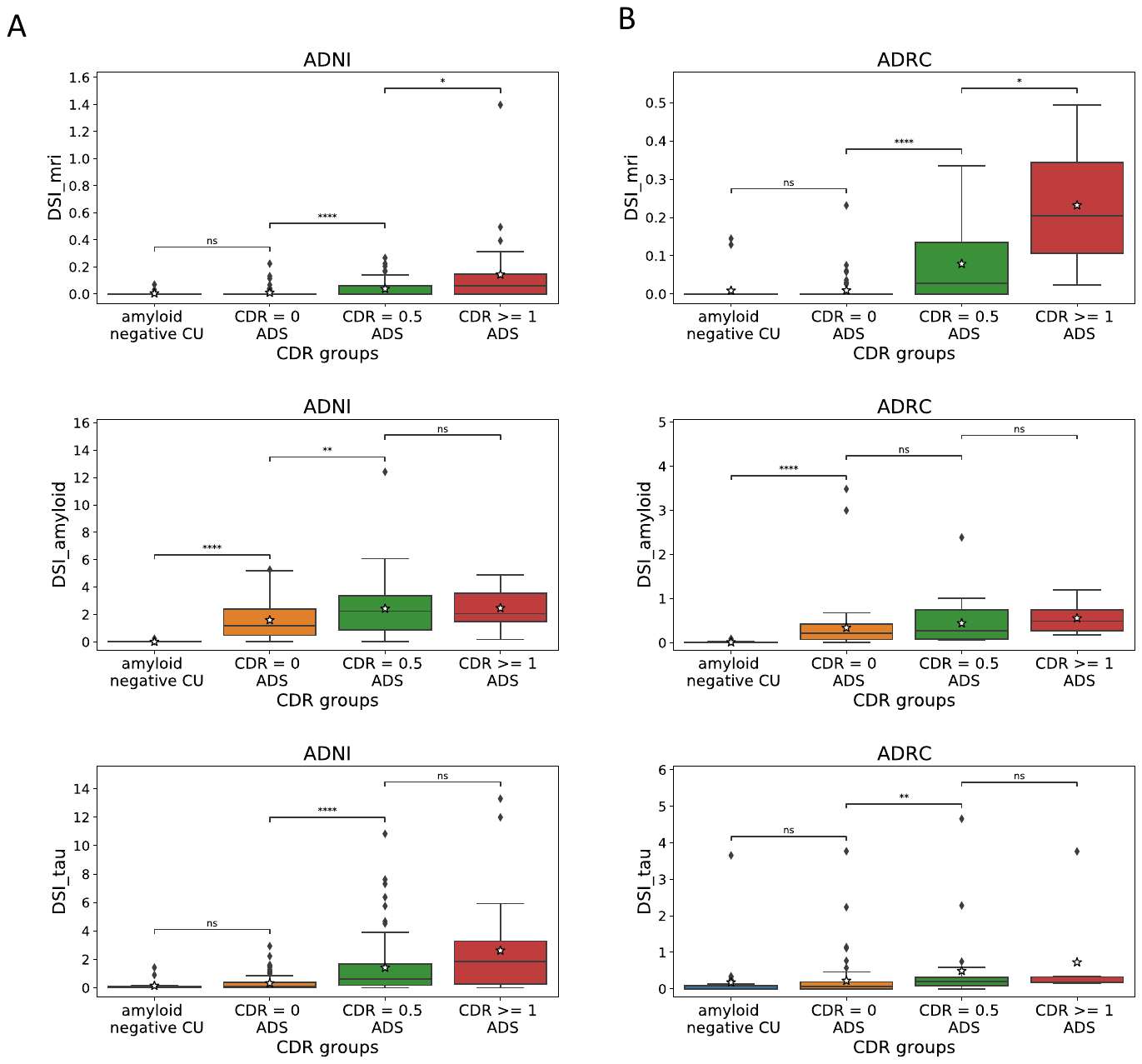

#### **Figure S3**: Box plot showing DSI across each modality for both ADNI (**S3A**) and Knight ADRC (**S3B**). The figures from top to bottom represent DSI_mri, DSI_amyloid and DSI_tau respectively. The x-axis shows the different CDR groups in the ADS and CU-test (Section 2.2.1 and 2.3.1). FDR-corrected post hoc Tukey comparisons used to assess pairwise group differences. Abbreviations: DSI: Disease Severity Index, CDR = Clinical Dementia Rating. Statistical annotations: ns: not significant 0.05 < p <= 1, * 0.01 < p <= 0.05, ** 0.001 < p < 0.01, *** p < 0.001.
